# Modelling movement and landscape connectivity of New Zealand native birds in highly structured agroecosystem networks

**DOI:** 10.1101/2020.06.24.170274

**Authors:** Jingjing Zhang, Jennifer L. Pannell, Bradley S. Case, Graham Hinchliffe, Margaret C. Stanley, Hannah L. Buckley

## Abstract

Understanding how spatial heterogeneity affects movement and dispersal is critical for maintaining functional connectivity in agroecosystems. Least-cost path models are popular conservation tools to quantify the cost of a species dispersing though the landscapes. However, the variability of species in life history traits and landscape configurations can affect their space-use patterns and should be considered in agroecosystem management aiming to improve functional biodiversity. In this study, we modelled the connectivity properties of native species on a real agroecosystem landscape dominated by sheep and beef farming in north Canterbury, New Zealand, where the recovery of native bird population is desired. We chose two species to act as case studies that were contrasting in their mobility: New Zealand pigeon/kererū (*Hemiphaga novaeseelandiae*; highly mobile) and southern brown kiwi/tokoeka (*Apteryx australis*; flightless). Networks of the least-cost paths of kererū and tokoeka were constructed based on their habitat preferences and movement capacities, and we compared and contrasted the connectivity properties and network topographies of their networks. We then compared the network metrics of western side (higher density of forest) with the eastern side (dominated by grazed grassland) of the study area where the vegetation composition was vastly different for both species. The results shown three variables were the most important contributors to the structure of the dispersal networks: the nature of the matrix, spatial structure of vegetation patches, and the gap-crossing ability of the study species. Tokoeka were able to utilise smaller habitat patches as stepping-stones for dispersal, while kererū can select more preferred habitat patches due to their high movement capacity. In contrast to the eastern side, we observed the western/forested side to have more, and stronger, links among habitat patches for both species, due to the presence of several large patches of native forest. Our work suggested that one size does not fit all, rather, conservation strategies that account for species’ life histories and movement traits are required to identify and preserve a connected ecological network.

## Introduction

Agricultural intensification leads to loss and degradation of natural habitat, and alterations in resource availability for native animals (Estrada et al., 2012; Fahrig et al., 2011). Habitat destruction typically leads to fragmentation, the division of habitat into smaller and more isolated fragments separated by a matrix of human-transformed land cover. Such changes undermine the ability of many organisms to move throughout landscapes (Fischer & Lindenmayer, 2007). Reducing the movements of individuals restricts their ability to disperse, recolonise and occupy new habitat patches, and thus fragmentation poses a significant threat to the viability of populations (Damschen et al., 2006; Harris & Silva-Lopez, 1992). Agroecosystems are often dominated by generalist species that can survive in fragmented habitats and are tolerant of human and livestock disturbances (Felton et al., 2010; Norton & Pannell, 2018). Agroecological approaches have been proposed as ways to promote biodiversity and mitigate the detrimental effects of agriculture on native species (Altieri, 1999; Tscharntke et al., 2012) (Kremen et al., 2012); however, such practices often require specific knowledge of the interactions between biotic and abiotic agroecosystem components (Duru et al., 2015). Thus, it is important to understand and quantify how animals response to the different habitat configurations that can result from agriculture, and, in particular, how landscape structure affects movements and dispersal (Adriaensen et al., 2003; Swihart et al., 2003).

Landscape connectivity, the degree to which structural landscape elements facilitate or inhibit movement between locations (also referred as ‘functional landscape connectivity’), is a widely used concept to improve conservation management and planning (Bélisle, 2005; Fischer & Lindenmayer, 2007). Connectivity is broadly applied as a way to prioritise areas for restoration and protection (e.g. Fuller et al. (2006)), assess the placement and effectiveness of habitat corridors (e.g. Pardini et al. (2005)) and predict the spread of invasive species (e.g. Stewart □ Koster et al. (2015)). Enhancing the ability of individuals to move through a landscape has therefore become a priority in many conservation strategies because it supports population growth and mitigates risk of local extinctions in isolated sub-populations (Crooks & Sanjayan, 2006). More recently, it has become a focus in agroecology (Dondina et al., 2018). In practice, identification of existing corridors and identification of where restoration of permeable habitat is needed are the first steps in promoting conservation of native species within a given agroecosystem (Clauzel et al., 2015). Approximately 40% of New Zealand’s land area is used for sheep and beef farming (Norton & Pannell, 2018), and remnant forest patches are common within sheep and beef farms. Addressing knowledge gaps on how native species with different behavioural characteristics interact with fragmented landscape in agroecosystems, especially when sharing with human and other species, is a vital staring point of improve connectivity for conservation in these systems (Norbury et al., 2013).

Methods to identify landscape connectivity elements and assess their role in population dynamics involve network modelling, and a variety of algorithms have been developed for this purpose, such as least-cost paths modelling and circuit theory (Avon & Bergès, 2016; Howey, 2011; LaPoint et al., 2013). Both methods have been broadly used in conservation (Avon & Bergès, 2016; LaRue & Nielsen, 2008; Yumnam et al., 2014), and a recent study evaluated the common landscape connectivity metrics using spatially-explicit simulation found least-cost path modelling out-performs circuit theory in the strength of correlation with true connectivity (Simpkins et al., 2018). Least-cost path model calculates routes of maximum efficiency (i.e., ‘lowest cost’) between two points, assuming that an individual has knowledge of the composition and configuration of the landscape (Adriaensen et al., 2003; Sawyer et al., 2011). There are two main input components to a connectivity model: the nodes, and a cost-surface or resistance surface (Newman, 2003). The nodes represent locations that animal may travel from and to, and traditionally treated as spatial points of the centroids of the habitat patches (Galpern et al., 2011). Cost-surfaces are raster representations of landscapes that describe the degree of difficulty an individual experiences in traversing a grid cell. The cost value assigned to each cell is based on both landscape elements or features, and species-specific factors that influence movement (e.g. mortality risk, energy expenditure, or behavioural aversion), with low cost values indicating higher willingness to cross the landscape and vice versa. The cost of effect distance on a landscape is represented by the resistance distance between the nodes in the network.

There are numerous metrics for condensing information from networks (Costa et al., 2008). Agricultural areas differ from natural ecosystem in vegetation type and intensity of production system, so they represent an extreme condition in ecological networks (Gliessman, 2014). Metrics that are more relevant to agroecosystem include those describe network topology, such as the cumulative frequency distribution of the links per node (or degree distribution) and the level of connection among the nodes (or connectance), which allow us to assess the functional overlap of species. The network-wise properties, including the similarities in the values between connected nodes (or assortativity), and the overall clustering coefficient between nodes can also be measured (Bohan et al., 2013).

This study aimed to quantify the effects of different life history trails such as movement and habitat preference on the dispersal network of native birds in agroecosystems. To demonstrate the effect of movement ability on functional connectivity, we applied the least-cost path analysis on two native birds as examples, which are markedly different in their movement and foraging traits. The species were selected so that they represented birds at either end of the mobility spectrum, with kererū (*Hemiphaga novaeseelandiae*) as a generalist frugivore with high mobility, and southern brown kiwi (*Apteryx australis*) as a flightless insectivore. we applied a least-cost path approach to compare functional corridors of the two species explicitly on a real agroecosystem landscape, where the detailed vegetation information is available (Norton & Pannell, 2018). To access the influence of landscape structures and vegetation types on the connectivity properties, we also compared the modelling results of the study species between the eastern (dominated by forest) and western (dominated by pasture) sides of the study area.

## Methods

### Study area

The *c*. 18 km^2^ study area landscape in North Canterbury, New Zealand, comprised a pastoral farming landscape dominated by sheep and beef farmland intermixed with other land uses including exotic forestry plantations, patches of remnant and regenerating native tree and shrub vegetation, exotic shrub patches, and small areas of dairy farming and horticulture. The shrubby vegetation was mostly continuous kānuka (*Kunzea* spp.), matagouri (*Discaria toumatou*), and gorse (*Ulex europaeus*). This vegetation mixture, typical of New Zealand South Island sheep and beef farming landscapes (Norton & Pannell, 2018), is highly fragmented and contains little remnant or old growth forest; however, larger forest patches were present predominantly in the south-western corner of the study area, including a pine (*Pinus radiata*) plantation (1,630 ha), some continuous mixed native forest patches (250 ha), and an old growth native forest patch (110 ha). Such remnant forest patches have potential to provide resources and habitat for a range of native bird species, particularly where invasive mammalian pest control is occurring.

#### Reclassified landscape

To create vegetation coverage maps that are relevant to the habitat quality of the study species, we combined landscape information from a thematic classification of 30-cm resolution RGB aerial imagery and three spatial land cover/land use databases: the New Zealand Land Cover Database (LCBD, version 4.1), the Land Use Carbon Assessment Survey dataset (LUCAS) and the Agribase farm property and land use dataset. The original categories from each data source were reclassified, as indicated table A1 in Appendix 1, and were re-defined by the vegetation types from automatic classification of aerial photographs (Figure 1). The RGB aerial imagery was classified into dominant woody vegetation types and canopy density classes using a semi-automated, object-based classification procedure within the e-Cognition^®^ image segmentation and classification software. The supervised image classification algorithm was trained by one of the authors using vegetation types identified at a set of 500 random points across the image and confirmed by an expert ecologist with extensive knowledge of the vegetation types in the region. The woody vegetation objects (polygons) were classified as: mixed native broadleaf, kānuka, matagouri shrubland, native old growth remnant vegetation, exotic deciduous, exotic conifer, and gorse shrubland. Functionality within the eCognition software also enabled us to categorise vegetation segments of each type into three density classes: continuous forest (>70% forest cover), diffuse forest cover (15-70% cover), and sparse forest (<15% cover). Overall, a classification accuracy of 70% was achieved across all vegetation types, quantified by comparing mapped vegetation types against the true vegetation types at 100 additional locations. This comprehensive set of spatial land cover/land use data from different sources, in combination with spatial information regarding the locations of roads, enabled the development of a final landscape layer depicting features affecting resource provision and movement potential for avian species in this study landscape.

**Figure 1.**
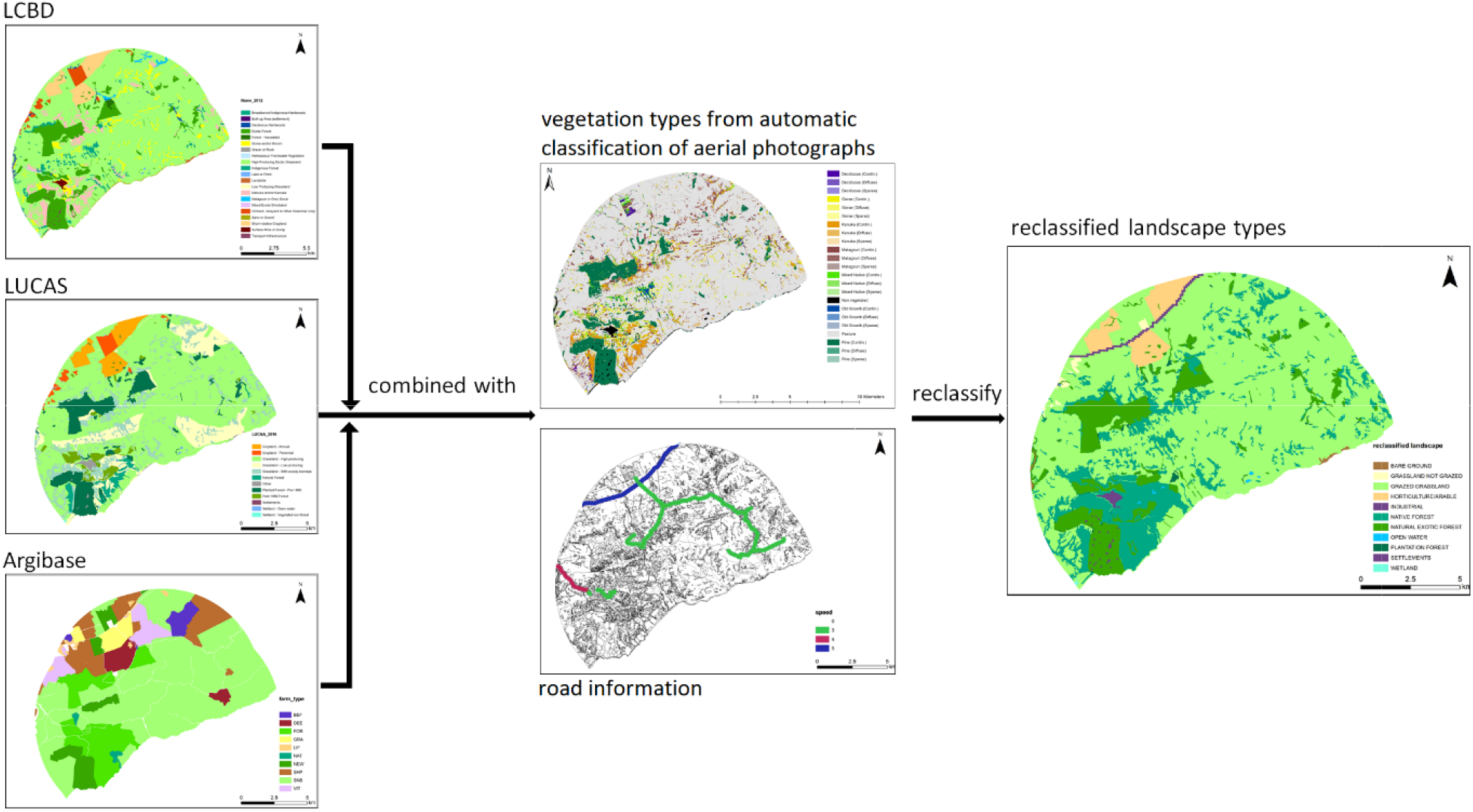
Workflow for generating the reclassified landscape from various spatial datasets.

### Focal species and habitat

kererū are primarily frugivorous and have a broad diet that includes both native and introduced plant species (Clout & Hay, 1989; Higgins & Davies, 1996). They are capable of making long distance flights of up to 18 kilometres to take advantage of seasonal food sources, but mainly occupy an areas with limited movements over several weeks (Baranyovits, 2017; Clout et al., 1991; Wotton & Kelly, 2012). As a contrast, we chose flightless southern brown kiwi/ tokoeka to represent the second type of native bird. The tokoeka are distributed mainly within indigenous forests of Haast, Fiordland and Stewart Island (Robertson 2007). There is limited information about this critically endangered sub-species of brown kiwi, except for a few genetic and breeding ecology studies (Colbourne, 2002; Herbert & Daugherty, 2002). We used the home range and dispersal distance information of similar species the northern brown kiwi (*A. mantelli*), which was considered conspecific with the tokoeka until 2000 (Burbidge et al., 2003). The home range of northern brown kiwi has been estimated at 2.03 ha with a mean maximum dispersal distance of 337 meters between forest remnants (Potter, 1990).

Habitat patch suitability maps of the two example species were constructed based on the habitat preferences and foraging ecologies of the two species, following the recommendations of the Corridor Design Project (http://corridordesign.org) (Beier et al., 2008). Suitability is a unitless variable specific to the species scaling from 0 - 100 with the following breaks: 0 no use at all; 1 - 30 avoided; 30 - 60 occasional use for non-breeding; 60 - 80 consistent use for breeding; 80 - 100 best habitat for survival and breeding, see table A2 in Appendix 1 for detail. Based on literature and expert advice, a score was assigned to each cover type of the reclassified landscape (Table A2 in Appendix 1) to generate habitat maps for the scenarios representing the ecology of the two birds with high (kererū) and low (brown kiwi) movement capacities in this landscape (Cunningham & Castro, 2011; Elliott et al., 2010; Potter, 1990; Powlesland et al., 2011; H. A. Robertson et al., 2013). Roads were considered as unsuitable and given a zero value, and the quality score of any habitat within 150 m of road was reduced by 10%. Furthermore, habitat patches that were smaller than minimum home range size (20 ha for kererū and 2.03 ha for brown kiwi) or too isolated (more than 4620 m from nearest patch, maximum pasture crossing distance for kererū (Pierce & Graham, 1995), and more than 337 m from nearest patch for tokoeka (Potter, 1990)) to be used by the species were reduced in suitability by 50%. Finally, all areas with a habitat quality score higher than 60 were defined as habitat. Because the scores of habitat types were given based on species-specific preference of land-use, the habitat maps of the two species also reflected such species-specific variations (Figure 2).

**Figure 2.**
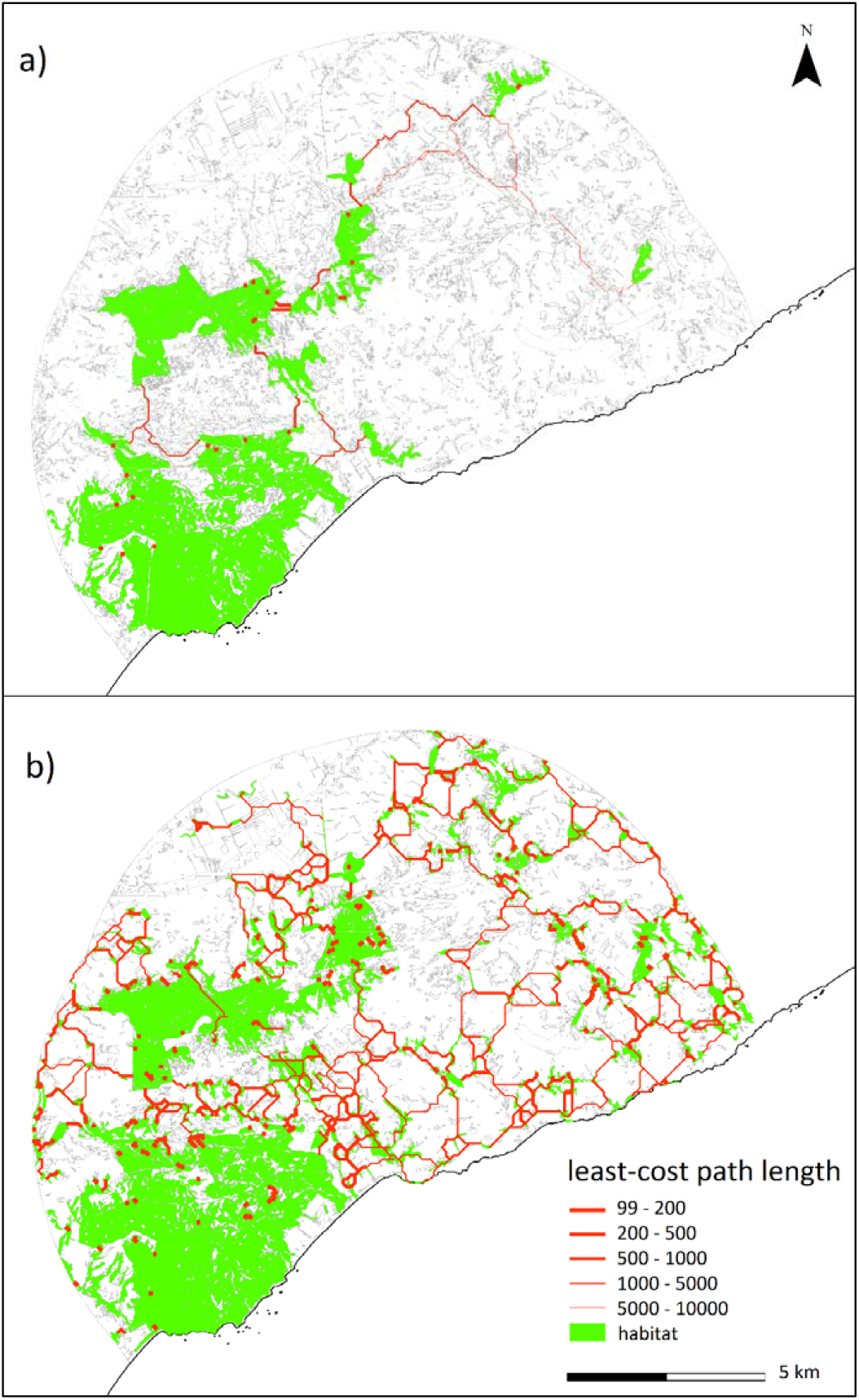
Least-cost paths of a) the mobile species (kererū) and b) the flightless species (tokoeka) as the model outputs. The grey lines indicate the outline of the study area and polygons of landscape patches within the area. The black line represents the coastline.

### Resistant surface

The vegetation type maps were converted into raster surfaces with a cell-size of 100 m. Species-specific maps of dispersal resistance were established in the non-habitat area. Each raster cell in the resistance map was provided with a value that marked how difficult it was to move across the cell (Poor et al., 2012; Sawyer et al., 2011; Zeller et al., 2012). Resistance values were assigned following a scale that doubles between classes (from 1 - 32 in geometric intervals), based on species knowledge about willingness to cross a vegetation type, physiological cost of movement across a vegetation type, and survival rate in a vegetation type (see table A3 in Appendix 1 for detail). For example, kererū may be equally willing to cross pasture with shrubs as empty fields, but more resources and perching places in shrubs and possibly lower likelihood of predation by raptors reduce cost and increase survival (Campbell, 2006). Similarly, roads and settlements are considered a cause of death for native birds, and were given the highest resistance values (Prendergast et al., 2006; 2013).

### Network model and analysing connectivity

We performed least-cost path connectivity analysis based on the species-specific habitat patches and resistant surfaces using the Linkage Mapper software (McRae & Kavanagh, 2011) for each of the two species. The aim of the network analysis is to calculate the least cost paths across resistance surface between all habitat nodes (the habitat polygons in our case) within maximum gap crossing distance of species, then use the least cost paths as network edges for descriptive statistics. Habitat networks were assembled by connecting habitat patches (i.e. nodes) form edge to edge via least-cost paths (i.e. links) through resistance surfaces (Adriaensen et al., 2003). We employed the Linkage Mapper programme to calculate the effective distances between the habitat nodes: First, Linkage Mapper found adjacent core areas and created a network of core areas using adjacency and distance data. Second, it calculated cost-weighted distance and least-cost paths; finally, it measured least-cost corridors and used them to create a single map. Several measurements were then carried out to describe network topologies (see Newman (2003). We first assessed the total corridor numbers between the two species, and for each species, the corridor numbers and link density per node. Second, to quantify the robustness of the network to node detentions, we also calculated and compared the degree distribution and connectance, which can help to identify highly connected or potential ‘keystone’ habitat patches or groups (Ledger et al., 2013). Third, correlation between the directed degree and undirected degree of habitat patches with non-zero degrees were measured, which gives an indication of whether some patches acted as steppingstones for movement. Finally, the cluster coefficients were evaluated, which measure the local group cohesiveness, and give an indication of how frequent the chance of animal to move between habitat patches. Because the western and eastern sides of the study area demonstrated distinct feather in the distribution and composition of vegetation patches, we also compared the connectivity features between the two sides for both species. Models with maps were developed in ArcMap v. 10.6 (Environmental Systems Research Institute, 2017) and networks were created using the Linkage Mapper toolbox (McRae & Kavanagh, 2011). Network properties analysis was carried out in R (R Core R Core Team, 2019) using ‘network’ (Butts, 2008), ‘igraph’ (Csardi & Nepusz, 2006) and NetIndices’ (Soetaert et al., 2015) packages.

## Results

### Landscape structure and habitat networks

The habitat networks of the two example bird species exhibited distinct features. For kererū, there were 25 core habitats identified, with a total area of 2876.92 ha. There were a higher number of habitat patches identified as available for tokoeka, and the 420 habitat patches made up a total area of 4374.69 ha. The habitat patches of kererū comprised 16.64 % of the total area of the study site and has a mean area of 110.65 ± 38.81 ha (mean ± 1 *SE*), while the tokoeka habitat, although constituted 25.3 % of the total area, were smaller on average (10.32 ± 2.65 ha).

The habitat patches distribution features also differed between the western and eastern sides. For both species, there were more areas that can be used as their habitat on the western side due to a large and continuous patch of native forest (26.51 % of the total area with a mean area of 125.26 ± 45.30 ha per patch for kererū, and 35.08 % of the total area with a mean area of 14.19 ± 4.36 ha per patch for the tokoeka). On the eastern side the landscape was dominated by grazed grassland, and the suitable habitat patches were sparser and with a smaller mean area for both kererū (30.27 ± 4.91 ha) and tokoeka (4.36 ± 0.44 ha).

### Least-cost paths

Linkage Mapper created corridors for both animals, and the networks of the two bird species show divergent connectivity properties. For kererū, there were 31 links/least-cost paths identified and the mean distance for them was 1227.13 ± 220.47 m. Comparatively, there were more least-cost paths identified for tokoeka, 599 least-cost paths, with a shorter mean length of 390.62 ± 11.90 m.

For each species, the connectivity patterns also varied between the western and eastern sides. The number and mean length of the least-cost paths for kererū differed drastically between two sides. There were 26 links/least-cost paths identified and the mean distance for them was 515.72 ± 153.26 m on the western side, and only 4 long least-cost paths were identified on the eastern side with mean length of 6688.75 ± 2322.65 m. In contrast, as tokoeka were able to use smaller vegetation patches than kererū, many more least-cost paths were identified by the model on both sides of the study area; there were 343 links on the western side and 144 links on the eastern side, and their mean distances were much closer at 286.27 ± 21.96 m and 320.82 ± 23.16 m, respectively.

### Network topology comparison between kererūl and tekoeka

For both bird species, the degree distributions roughly followed a power rule whereby most nodes had less than six links, with a mean of 5.05 ± 0.31 for kererū and 3.13 ± 0.13 for tokoeka. The link density of the kererū network (2.72) was lower compared to tokoeka (3.23). For the tokoeka network, there were two important habitat nodes that generated 25 and 35 links on the western side. This indicated such a network was robust to losing random nodes but very vulnerable to the loss of just one or a few key nodes. The kererū network was simpler, with fewer nodes and links generated from them (less than seven links). However, the habitat patches within the kererū network were more closely related and connected to each other, indicated by a higher connectance value of 0.11, while within the tokoeka network, habitat patches interacted between each other at a relative lower level of overall connectance (0.008). The clustering coefficient quantifies how well connected are the neighbours of the vertex in the network (Soffer & Vazquez, 2005). For kererū, the network cluster coefficient value was 0.263, indicating an individual could take relative shorter path to get from one node to the next, compared to that of tokoeka (0.223). This was further confirmed as the degree of separation or the average number of nodes between two nodes was 4.15 for the kererū network, and 11.18 for the tokoeka network.

### Network properties between the western and eastern sides of the Canterbury site

The network topology also varies between the two sides for the study sites for both species. The vegetation type composition of the western side of the Canterbury site is more complexed and contains large forest patches, compared to the eastern side which is dominated by pasture landscape. For both species, the network of the western side had more links compared to the eastern side (Fig. 3). The link densities of kererū were 2.64 on the western side and 2.00 on the eastern side. The link density of tokoeka was higher on both the western and eastern sides compared to those of the kererū, being 2.98 and 2.21, respectively. The kererū network had fewer nodes and links generated from them, and the interaction among the others were strong, which was shown by connectance values of 0.13 and 0.67 for the western and eastern sides. For the correlation between habitat patches, the kererū east and west networks, and the west brown kiwi network were dis-associative (i.e. the connection between any two of the habitat patches was weak), demonstrated by negative correlation values (−0.27 and −0.87 for kererū, and −0.13 for the tokoeka west). However, the correlation value of the network for tokoeka on the eastern side was 0.09, which suggested this network was associative, that is, some habitat patches can act as steppingstones linking multiple other patches to facilitate dispersal of this species. Western networks for both species had lower clustering coefficient values (0.22 for kererū and 0.19 for tokoeka) compared to those of the eastern sides (0.60 for kererū and 0.31 for brown kiwi), indicating there were higher proportion of disconnected nodes to their neighbours in both networks.

**Figure 3.**
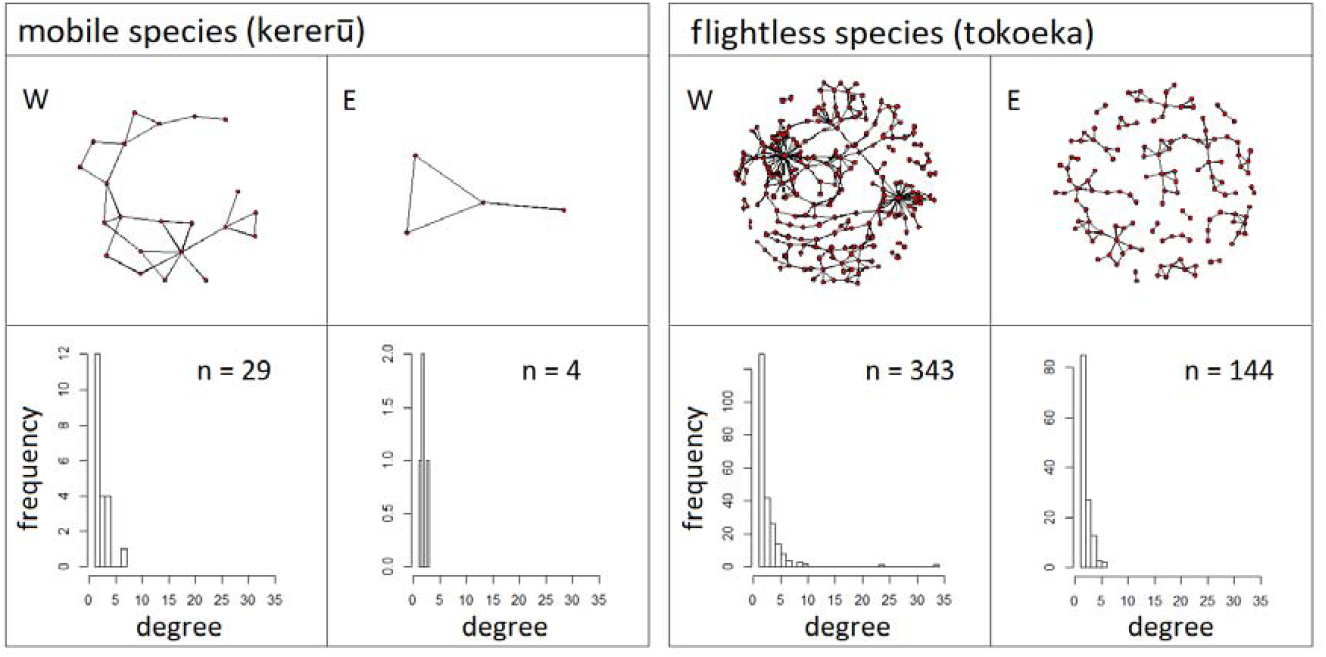
Networks of kererū and tokoeka shown in circular plots on the top panel, the figures labelled with ‘W’ represent the network structures of the western side and ‘E’ for the eastern side of the study site for each species. The red dots indicate nodes of habitat patches, and the black lines are links between nodes. Histograms of the degree distribution of each network are displayed in the bottom panel, where n is the total number of links.

## Discussion

Using a least-cost paths approach, we investigated the connectivity networks of two New Zealand native bird species, with different habitat preferences and movement capacities, within a typical agroecosystem landscape. Number and mean length of corridors reflected how the two species responded to the landscape structure differently in dispersal, and the different opportunities a landscape might provide for birds depending on their mobility. Our results suggested forest landscape patches were important in the network of both species, and were the keys to enhance the connectivity. In highly fragmented landscape, kererū can travel to habitat patches with features that they prefer. Meanwhile, in the same landscape, tokoeka least-cost paths had a mean distance close to its maximum dispersal range. For both species, the vegetation types and structure of the western side was more suitable due to the presence of large forest patches, and such key habitat patches offered many links to neighbouring patches for the bird to disperse. With the tokoeka, however, the network of eastern side was more vigorous compared to the western side, as the habitat patches were better correlated with each other and acted as steppingstones for the bird to commute between each other. Together, these findings show that native bird species with different habitat preference and movement capacities can have divergent space-use patterns within the same landscape, and that network modelling to identify the least-cost corridors and key features shaping the network is an effective first step towards locating key areas of functional overlaps of species, and conserving a connected ecological network in agricultural landscape.

The land-use patterns of two species generated from our model shown how species’ dispersal patterns may respond differently to the same physical landscape conditions. Such differences were influenced by three key factors: nature of matrix, spatial structure of vegetation patches, and the gap-crossing ability of the animal. Agroecosystems provide limited resource for the native animal species because of intensive land modification, which leads to the loss of habitat diversity and indigenous food sources (Altieri, 1999; Case et al., 2019). Compared to many introduced species, native birds in New Zealand rely more on indigenise forests for food and shelter (Montague-Drake et al., 2009). The matrix of vegetation in agroecosystems can provide novel resources for native vertebrates (e.g., exotic fruit trees as food sources), but these may not be adequate for providing the full range of resources required (Estrada et al., 2012; Stanley & Bassett, 2015). Our results shown large native forest patches acted as centroids of the dispersal networks in the spatial matrix of the landscape, suggesting that the existence of native forest was crucial to the survival and persistence of the study population in this agroecosystem. The difference in the nature of matrix between the western and eastern sides of the study site provided a convenient natural experiment that demonstrated the predominate predominant effect of the different combination of vegetations. Compared to the eastern side of the study site, the western side holds more and stronger links between the habitat patches for both species, due to the presence of large patches of native forest. This was especially important for the kererū as this species requires larger home range size across high-quality habitat (for fruit and nectar sources), compared to tokoeka that can occupy smaller habitat patches. Loss of large forest habitat significantly changes the possible network by reducing the possible paths of dispersal. This shows the vital importance of remnant and regenerating native forests in agroecosystems for enhancing biodiversity and reducing pressures from land-use and climatic changes on native species.

The spatial distribution of suitable habitat patches also played a significant role in shaping the network structure. For both species, suitable habitat patches were of higher density at the south-western corner of the study site, while at the entire eastern side they were sparse and disconnected. In this geographical context, colonization of new habitat patches can occur only if migration of individuals occurred in an easterly direction. For both species, there were asymmetrical connectivity between patches observed in the networks due to spatial distributions of the habitat patches. The dispersal of individuals between these pairs of nodes not necessary be balanced. The connectance results indicated that the least-cost paths or the corridors were clustered at certain area within the network, with the exemption of the eastern side of the study site in tokoeka’s network, where the chance of movement between patches can be more even out.

An individual’s gap-crossing ability was another key factor that interacted with landscape structure. Bird behaviour can be markedly differ under different landscape conditions (e.g. between fragmented agroecosystem and nature habitat) (Gallagher et al., 2017). In our model, kererū shown the ability to tolerate and used a wide range of forest and shrub landscapes, and can fly great distances largely undisturbed even with changes in matrix types. Because of their great gap-crossing ability and large home-range area, there might be little need to focus on improving connectivity at the scale of individual farms. Instead, planting the fruit trees on which they primarily feed would encourage individuals of this species to exploit and stay in an area. In contrast, tokoeka have very low gap-crossing ability, and enhancing connectivity through landscape corridors would be a priority, such as increasing the link density and network correlation coefficient. Compared to lowering dispersal costs of the landscape, more efficient methods to strength the tokoeka network might include creating islands of habitat to act as steppingstones, for example, increase the numbers of suitable habitat (i.e. small but larger than the home range of tokoeka) that are adjacent to each other that within the maximum dispersal distance of the animal.

Conserving landscape connectivity for multiple species is a preferred strategy for promoting biodiversity, and should be assessed by accounting for the needs of inhabited species that have divergent life histories and movement ecologies (Nicholson et al., 2006). Our study is the first to evaluate the similarities and differences of the connectivity properties of New Zealand native birds in agroecosystems. Specifically, the habitat patches were treated as nodes, rather than spatial points that were used in many other similar works studying invasive species dispersal. Our method improved the accuracy in corridor identification and is a more appropriate methods for measuring their utility (Galpern et al., 2011). This improvement in modelling technique allow the true costs of travelling between edges of habitat patches to be estimated Our results demonstrated the potential importance of large forest patches in retaining and promoting biodiversity, as these patches can be used as habitat for multiple species, as well as providing links to the other neighbouring patches in the dispersal networks.

Although the modelling results provides valuable insights, it is important to highlight that the landscape genetics approach we used to calibrate our habitat and cost-surface landscape representations is not the only or necessarily the best approach. Like many other cost-based corridor models, the habitat selection and resistance surface information is extracted from the literature (e.g. Huck et al. (2011); LaRue and Nielsen (2008)). Also, our model, as the other connectivity models, unrealistically assume an animal has complete knowledge of the landscape (Adriaensen et al., 2003). The validity of the model can be tested, which will allow the model to be improved, if the least-cost path predictions there were real movement trajectories of the study species to be compared to (as in Driezen et al., 2007; Poor et al., 2012; Walpole et al., 2012). In our case, there is limited data available on dispersal distances for these two species, particularly lacking in agroecological systems. Habitat suitability can be a poor proxy for landscape connectivity as animals are able to move through less suitable areas especially during long-distance dispersal events (Keeley et al., 2017). The networks of the two example species were deliberately chosen to represent native birds with different movement abilities, and modelled based on current empirical knowledge. Simulation models such as agent-based model can be employed to better understand how such life history traits affect connectivity, using individuals with various combination of movement capacities and habitat preferences.

## Conclusion

Our network approach identified and compared dispersal patterns of two bird species with differences in movement capability within a patchy agricultural landscape. Network topology measurements suggested that the spatial structure of the study area would be more hospitable to bird species that can tolerate lower quality habitat patches that are smaller in area. To maintain populations of birds with good gap-crossing ability, it is crucial to retain suitable high-quality habitat. This analysis illustrates that network analysis is a powerful tool for identification connectivity and can be used as a starting point for building a complex conservation management plan that accounts for differences in species’ life history traits.

## Acknowledgements

We thank David Norton for providing useful input regarding native bird species resource use and vegetation type distribution within the study landscape. This work was funded by the New Zealand Ministry of Business and Innovation/New Zealand Biological Heritage National Science Challenge grant number C09X1501.

## Appendix 1

**Table A1:** Landscape reclassification from LCBD, LUCAS, Agribase, and RGB aerial photo classification.

| Category | LCBD category | LUCAS category | Agribase category | Aerial photo |
| --- | --- | --- | --- | --- |
| Grassland with Trees | Exotic Forest |  |  |  |
| Grassland with Shrubs | Gorse and/or Broom, Matagouri or Grey Scrub | Grassland - High producing |  | Matagouri, Shrubland |
| Grassland not Grazed | Mixed Exotic Shrubland, Orchard, Vineyard or Other Perennial Crop |  | NAT, NOF | Gorse |
| Grazed Grassland | Low Producing Grassland, High Producing Exotic Grassland | Grassland - Low producing, Grassland - High producing | SHP, BEF, SNB, DEE, GRA, LIF, NEW, none |  |
| Plantation Forest | Exotic Forest, Forest - Harvested | Planted Forest - Pre-1990, Post 1989 Forest |  | Pine |
| Native Forest | Broadleaved Indigenous Hardwoods, Manuka and/or Kanuka, Indigenous Forest | Grassland - With woody biomass, Natural Forest | FOR | Old_Growth, Mixed Native Deciduous, Kanuka |
| Open Water | Lake or Pond | Wetland - Open water |  |  |
| Horticulture/Arable | Short-rotation Cropland | Cropland - Annual | HOR |  |
| Settlements | Built-up Area (settlement) | Settlements |  |  |
| Bare Ground | Gravel or Rock, Landslide |  |  |  |
| Natural Exotic Forest | Deciduous Hardwoods, Exotic Forest |  |  |  |
| Industrial | Surface Mine or Dump, Transport Infrastructure |  |  | Road |
| Wetland | Herbaceous Freshwater Vegetation | Wetland - Vegetated non forest |  |  |

**Table A2:** Suitability score given to each type of reclassified landscape categories, based on knowledge of the two example species.

| Category | Bird with high movement capacity (kererū) | Bird with low movement capacity (brown kiwi) |
| --- | --- | --- |
| Grassland with Trees | 60 | 60 |
| Grassland with Shrubs | 50 | 50 |
| Grassland not Grazed | 50 | 50 |
| Grazed Grassland | 20 | 20 |
| Plantation Forest | 70 | 70 |
| Native Forest | 100 | 100 |
| Open Water | 0 | 0 |
| Horticulture/Arable | 30 | 30 |
| Settlements | 30 | 30 |
| Bare Ground | 0 | 0 |
| Natural Exotic Forest | 80 | 80 |
| Industrial | 20 | 0 |
| Wetland | 30 | 30 |

**Table A3:** Resistant value given to each grid cell of raster surfaces depending on the reclassified landscape vegetation categories, based on knowledge of the two example species.

| Category | Bird with high movement capacity (kererū) | Bird with low movement capacity (brown kiwi) |
| --- | --- | --- |
| Grassland with Trees | 2 | 2 |
| Grassland with Shrubs | 8 | 2 |
| Grassland not Grazed | 8 | 8 |
| Grazed Grassland | 16 | 16 |
| Plantation Forest | 1 | 1 |
| Native Forest | 1 | 1 |
| Open Water | 2 | 32 |
| Horticulture/Arable | 2 | 16 |
| Settlements | 32 | 32 |
| Bare Ground | 2 | 16 |
| Natural Exotic Forest | 1 | 1 |
| Industrial | 32 | 32 |
| Wetland | 2 | 30 |

